# Are the neural correlates of consciousness in the front or in the back of the cerebral cortex? Clinical and neuroimaging evidence

**DOI:** 10.1101/118273

**Authors:** Melanie Boly, Marcello Massimini, Naotsugu Tsychiya, Bradley R. Postle, Christof Koch, Giulio Tononi

## Abstract

The role of the frontal cortex in consciousness remains a matter of debate. In this Perspective, we will critically review the clinical and neuroimaging evidence for the involvement of the front versus back of the cortex in specifying conscious contents and discuss promising research avenues.

---

Consciousness is subjective experience, the ‘what it is like’ to perceive a scene, recognize a face, hear a sound, or reflect on the experience itself (Tononi et al., 2016a). Identifying the neural correlates of consciousness is important scientifically and clinically, to improve the detection of awareness and to design new therapies in patients who remain unresponsive after brain damage (Gosseries et al., 2014). While frontal cortex is crucial for intelligent behavior and cognitive control, its involvement in consciousness remains a matter of debate (Koch et al., 2016b). It has been widely assumed that prefrontal circuits are essential for consciousness, either alone (Del Cul et al., 2009) or in conjunction with parietal areas (fronto-parietal network (Bor and Seth, 2012; Laureys and Schiff, 2012)). In this Perspective, we will critically review the evidence for the role of the ‘front’ versus the ‘back’ of the cortex in supporting consciousness. In the ‘front’ we include pre-rolandic neocortex in dorsolateral, medial prefrontal, anterior cingulate, and orbitofrontal areas. In the ‘back’ we include neocortical regions within the parietal, occipital and lateral temporal lobes. We will not review the role of the medial temporal lobe or insular cortex nor the complex relationships between long-term memory, interoception and consciousness (Craig, 2010; Seth et al., 2011; Quiroga, 2012). However, lesion data (Corkin, 2002; Damasio et al., 2013) suggest that none of these areas is necessary for conscious experiences. We will also not discuss the role of different brainstem and subcortical meso-circuit structures in regulating the level of consciousness (Brown et al., 2010), although lesion data suggest that after widespread cortico-thalamic damage, consciousness can be absent even when brainstem activity is preserved (as in patients in the vegetative state (VS)) (Laureys et al., 2004).

## I. Dissociating the neural correlates of consciousness from other neural processes

### I.1. Neural correlates of consciousness: definition

The neural correlates of consciousness (NCC) are defined as the minimal neural mechanisms jointly sufficient for any one conscious percept (Crick and Koch, 1990).

<u>Content-specific NCC</u> are the neural mechanisms specifying particular phenomenal contents within consciousness - such as colors, faces, places, or thoughts. Experimentally, content-specific NCC are typically investigated by comparing conditions where specific conscious contents are present versus absent. The <u>full NCC</u> is constituted by the union of all content-specific NCC (Koch et al., 2016b). Experimentally, the full NCC can be identified by comparing conditions where consciousness as a whole is present versus absent - such as dreaming versus dreamless sleep. In principle, the full NCC can also be approximated by sampling the wide range of content-specific NCC. In practice, these two approaches progress hand in hand (Boly et al., 2013).

### I.2. Distilling the ‘true NCC’

Recent results have stressed the importance of dissociating the true NCC from other neural processes (Miller, 2007; de Graaf et al., 2012; Tsuchiya et al., 2015) that can be considered as ‘prerequisites’ and ‘consequences’ of consciousness (Aru et al., 2012) or alternatively as preceding and following the experience itself (Pitts et al., 2014; Tsuchiya et al., 2016). However, processes such as selective attention (and thereby also working memory) and memory retrieval may not be part of the NCC without necessarily happening before or after but rather concurrently with experience (Postle, 2015; Tononi et al., 2016a). Here factors that modulate the NCC without being directly involved in specifying conscious contents will be called ‘background conditions’ (Koch et al., 2016b). These include global enabling factors (Fig. 1), such as neuromodulatory systems, which influence the <u>level of consciousness</u> by modulating the activity of large parts of the full NCC (de Graaf and Sack, 2014). Other factors such as selective attention or expectation may modulate the probability of some <u>conscious contents,</u> often in a task-dependent manner (Aru et al., 2012). Other task-related neural mechanisms that may correlate with the occurrence of an experience include verbal and motor reports (Tsuchiya et al., 2015).

**Fig. 1.**
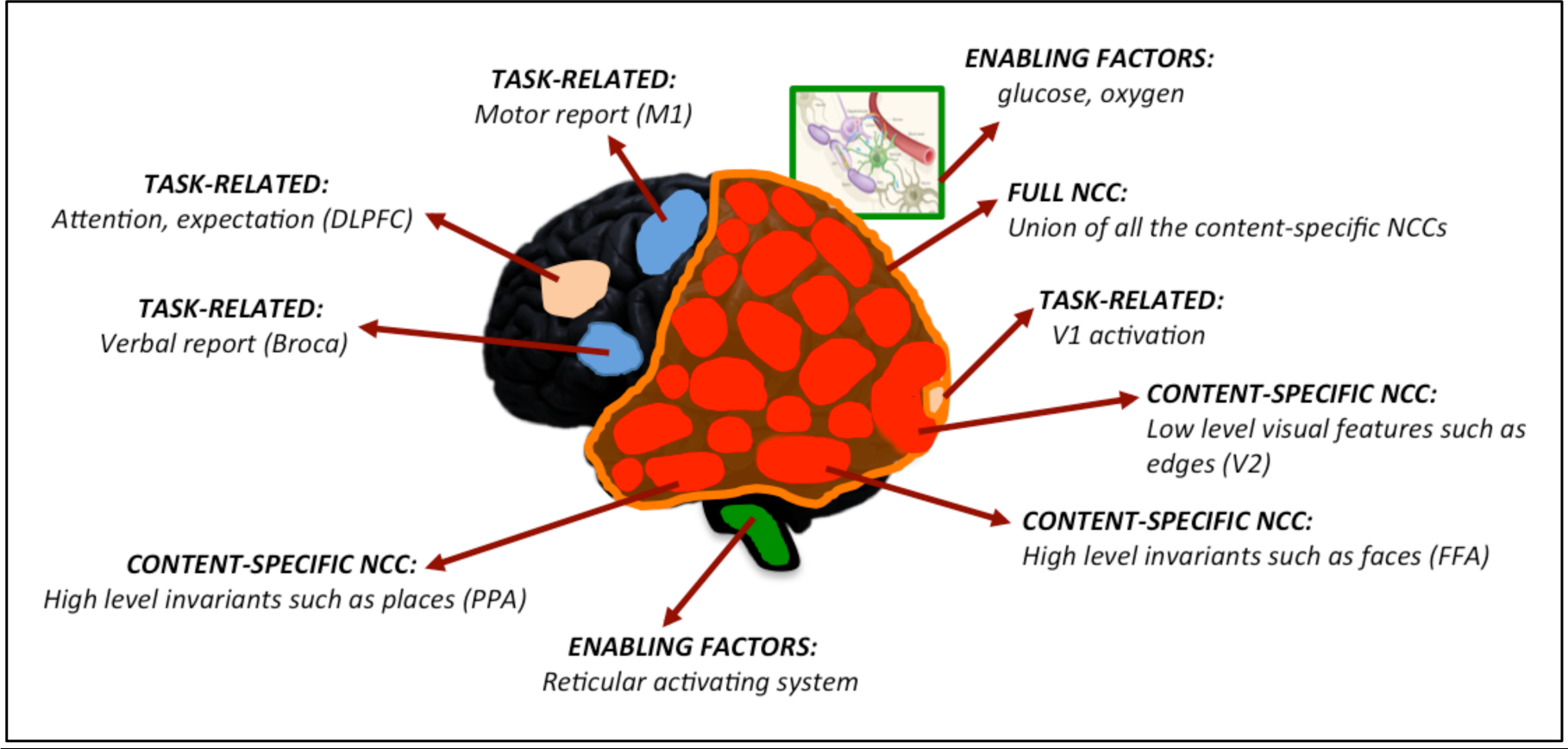
Definition of the neural correlates of consciousness (NCC). Content-specific NCC (in red) directly contribute phenomenal distinctions (such as low level visual features, faces or places) to consciousness. The full NCC (in orange) is constituted by the union of all the content-specific NCC. Background conditions can be divided into physiological or neural processes that enable or modulate the activity of the full NCC and thus influence the level of consciousness (enabling factors, in green) - such as brainstem arousal systems, glucose or oxygen - and task-related neural processes that modulate the activity of only some content-specific NCC (in beige) - such as attention, expectation, and possibly primary visual cortex (V1) activity. Other task-related neural processes such as verbal or motor reports (in blue) often follow the experience itself. DLPFC: dorsolateral prefrontal cortex; V2: secondary visual cortex; FFA: fusiform face area; PPA: para-hippocampal place area; Ml: primary motor cortex.

Several complementary methods can be used to distill the true NCC. First, with respect to the <u>full NCC, within-state paradigms</u> can be used to avoid confounds due to changes in behavioral state and task performance as well as to dissociate unconsciousness from unresponsiveness. For example, within either non-rapid eye movement (NREM) or REM sleep one can contrast neural activity when subjects report having dreamt (~60% of cases in NREM sleep, ~95% of cases in REM sleep) versus having been unconscious. Patients can also be conscious although unresponsive and disconnected from the environment in ~20% of cases during anesthesia (Sanders et al., 2012) and in ~35% of cases during complex partial seizures (Johanson et al., 2003). In such cases, methods assessing the complexity of neural EEG responses to transcranial magnetic stimulation (TMS) can also assess the presence versus absence of consciousness in unresponsive subjects (Casarotto et al., 2016). Second, with respect to <u>content-specific NCC,</u> experiments can be carefully designed to systematically investigate possible <u>dissociations</u> between consciousness and various cognitive functions such as report or selective attention (Aru et al., 2012; Koch and Tsuchiya, 2012). Third, it is important to quantify the systematic <u>associations</u> between the activity of a candidate NCC and the presence of a particular conscious content systematically across a large number of different experiments (Crick and Koch, 1990). Fourth, machine learning approaches can help to identify the true NCC as the neural activity patterns with the highest <u>predictive value</u> for specific conscious percepts (Sandberg et al., 2014). In practice, the true NCC should remain lawful across all dissociation, association and prediction approaches applied within lesion, stimulation and recording studies (Koch et al., 2016b). With these methodological clarifications at hand, we will now critically review evidence for the NCC in the front versus the back of the cerebral cortex.

## II. Clinical evidence for a contribution of anterior versus posterior cortex to consciousness

### II.1 Lesions

Lesion data offer strong causal evidence for the involvement or lack of involvement of different brain areas in supporting consciousness and its contents (Farah, 2004). Several well-documented patients have been described with a normal level of consciousness after extensive frontal damage. For example, patient A. ((Brickner, 1952), Fig. 2A), after complete bilateral frontal lobectomy, (*sic*) ‘toured the Neurological Institute in a party of five, two of whom were distinguished neurologists, and none of them noticed anything unusual until their attention was especially called to A. after the passage of more than an hour’. Patient K.M. (Hebb and Penfield, 1940) had a near-total bilateral frontal resection for epilepsy surgery, after which his IQ improved. Patients undergoing bilateral resection of prefrontal cortical areas for psychosurgery (Mettler et al., 1949), for example, Brodmann areas 10, 11, 45, 46, 47 or 8, 9, 10 or 44, 45, 46, 10 or area 24 (ventral anterior cingulate), remained fully conscious (see also (Penfield and Jasper, 1954; Kozuch, 2014; Tononi et al., 2016b)). A young man who had fallen on an iron spike that completely penetrated through both of his frontal lobes went on to marry, raise two children, have a professional life, and never complained of loss of sensory perception or other deficits (Mataro et al., 2001). A young woman with massive bilateral prefrontal damage of unclear etiology (Markowitsch and Kessler, 2000), had deficits in cognitive functions supported by the frontal lobe, but her consciousness and perceptual abilities were preserved. Medial prefrontal lesions, especially those involving anterior cingulate cortex, can cause akinetic mutism (Cairns et al., 1941), where patients visually track examiners but do not respond to command. Patients who recover from this state typically report that they were conscious but lacked the motivation to respond (Damasio and Van Hoesen, 1983).

**Fig. 2.**
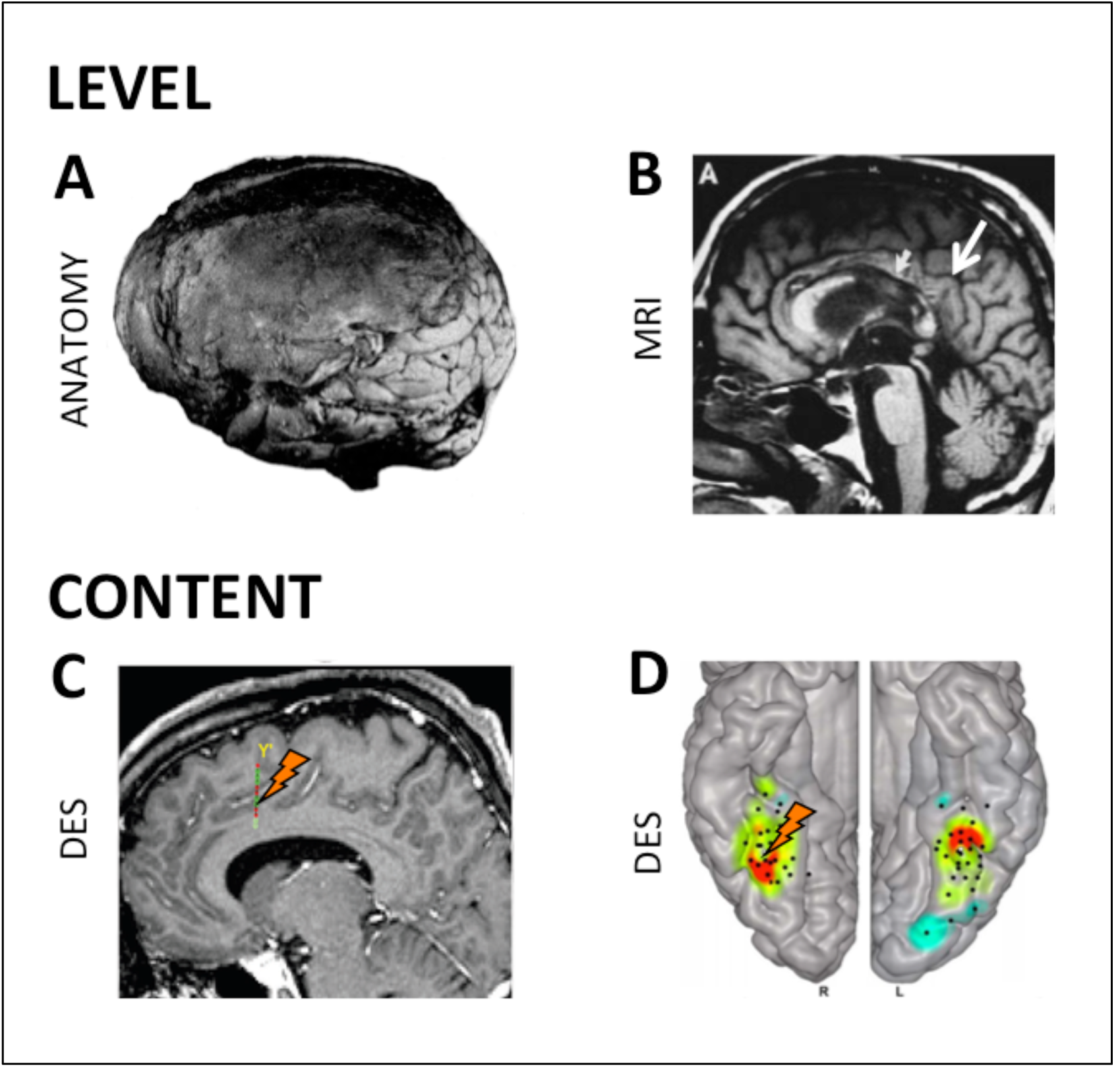
Clinical evidence. Upper panel: **(A)** Complete bilateral frontal lobectomy does not noticeable change the level of consciousness or social interactions (Brickner, 1952). **(B)** Lesions of posterior corpus callosum predict permanent vegetative state (Kampfl et al., 1998). **(C)** A recent study suggests that intrusive-thoughts can be elicited by electrical stimulation of anterior cingulate cortex (Popa et al., 2016). Eliciting any experience is, however, far more common when stimulating posterior than anterior cortical structures (Selimbeyoglu and Parvizi, 2010). **(D)** Direct electrical brain stimulation indeed supports a causal role for different parts of the posterior cortex in specifying conscious content, such as here for the right fusiform face area (FFA) in contributing to face percepts (Rangarajan et al., 2014).

Moving to the back of the brain, broad bilateral lesions (or ablations) of posterior cortex are extremely rare (Cavanna and Trimble, 2006). However, traumatic lesions of the posterior corpus callosum, connecting large parts of the posterior cortex, are found in 98% of patients who remain in VS after one year ((Kampfl et al., 1998), Fig. 2B). Moreover, such lesions are associated with a 214-fold risk of permanent VS (Kampfl et al., 1998). Posterior corpus callosum lesions also predict permanent coma after cardiac arrest (Bianchi and Sims, 2008). In contrast, while traumatic lesions of the frontal lobe are found in about half of patients with traumatic VS in the acute phase, they do not predict outcome (Kampfl et al., 1998).

There is abundant neurological evidence that lesions in the posterior cortex can cause a loss of specific contents of experience (Farah, 2004). For example, lesions of the right fusiform face area (FFA) cause prosopagnosia, an inability to recognize faces (Barton and Cherkasova, 2003). Lesions of infero-lateral occipital cortex cause achromatopsia, an inability to distinguish colors (Barton, 2011), which, when severe, may be accompanied by unawareness of the deficit (von Arx et al., 2010). In turn, other lesions of the occipital cortex lead to visual form agnosia, a selective inability to identify objects, or simultanagnosia, an inability to perceive more than one object at a time (Farah, 2004). Lesions of the postrolandic cortex lead to a loss of somatosensory percepts, lesions of left and right angular gyrus impair understanding of speech and prosody, respectively (George et al., 1996), and lesions of the inferior parietal lobule cause a loss of motor awareness (Sirigu et al., 2004). Lesions in left lateral temporal cortex may also lead to selective deficits for perception of single words or full sentences (Blumenfeld, 2011). By contrast, there is little evidence for loss of specific conscious contents after frontal damage (Penfield and Jasper, 1954). For example, lesions of Broca’s area, while impairing speech production, do not typically cause any loss of conscious speech perception (Blumenfeld, 2011). While frontal injuries can slightly increase the threshold for perceiving some brief (16ms) and masked visual stimuli, patients still experience them (Del Cul et al., 2009) - suggesting that these frontal regions may modulate the NCC rather than contributing directly to them (Kozuch, 2014).

### II.2 Stimulation studies

Electrical stimulation during neurosurgery is another important source of evidence for a direct contribution of different brain areas to consciousness (Penfield, 1959; Desmurget et al., 2013), as indicated by its superior value in predicting postoperative deficits compare to functional MRI (fMRI) or diffusion tensor imaging (Borchers et al., 2011).

It was recognized long ago that electrical stimulation of most of the frontal cortex has no consequence on experience (Penfield and Jasper, 1954) although it can interfere with tasks as well as induce involuntary movements (Selimbeyoglu and Parvizi, 2010). Transcranial magnetic stimulation (TMS) of the frontal cortex also does not seem to modify experiential content although it can interfere with speech production (Pascual-Leone et al., 1991)). Complex hallucinations have been occasionally reported after stimulation of the middle and inferior frontal gyrus (Blanke et al., 2000), similar to those classically reported after stimulation of temporal and parahippocampal regions (Penfield and Jasper, 1954; Megevand et al., 2014), suggesting a network effect. Recently, two case report studies however described the occurrence of a will to persevere and of intrusive thoughts after stimulation of the anterior cingulate cortex (Parvizi et al., 2013) and of mid-cingulate cortex ((Popa et al., 2016), Fig. 2C), respectively.

In contrast, electrical stimulation of the posterior cortex reliably induces discrete changes in the contents of consciousness (Selimbeyoglu and Parvizi, 2010). For example, direct electrical stimulation of early visual areas induces simple visual experiences such as phosphenes (Beauchamp et al., 2012; Winawer and Parvizi, 2016), which can also be induced by TMS of occipital and parietal cortices (Samaha et al., 2017). Electrical stimulation of postcentral gyrus induces somatosensory percepts (Penfield and Jasper, 1954), stimulation of temporo-parietal cortex induces experiences of visual motion (Rauschecker et al., 2011), while stimulation of right fusiform gyrus selectively disrupts the perception of faces ((Rangarajan et al., 2014), Fig. 2D). Moreover, the feeling of intention has been elicited in temporo-parietal cortex (Desmurget et al., 2009). Altogether, stimulation studies show consistent evidence for many conscious contents being specified by the posterior cortex, while the evidence is much less clear for the frontal cortex.

## III. Neuroimaging evidence for a contribution of the anterior versus posterior cortex to consciousness

Compared to lesions or electrical stimulations, neuroimaging studies offer less direct evidence for the contribution of any one brain region to consciousness (Farah, 2004). Indeed, functional activation maps frequently encompass brain areas that may not be directly involved in the mental process under investigation (Silvanto and Pascual-Leone, 2012; de Graaf and Sack, 2014). For example, while fMRI and intracranial EEG both reveal the activation of widespread bilateral temporo-occipital areas (beyond the FFA) after the presentation of faces, only in the right FFA does direct electrical stimulation disrupt face perception ((Rangarajan et al., 2014), Fig. 2D).

However, unlike clinical approaches, neuroimaging experiments can sample brain activity non-invasively in healthy volunteers (Poldrack and Farah, 2015). Importantly, they can also provide valuable information about the functional specificity of brain regions, if appropriate methodological steps are taken (Moran and Zaki, 2013; Poldrack and Farah, 2015). For example, neuroimaging experiments can demonstrate <u>dissociations</u> between the neural correlates of consciousness and those of other cognitive processes (Aru et al., 2012; de Graaf et al., 2012) by relying on <u>forward inference</u> (Henson, 2006; Moran and Zaki, 2013). Moreover, ever-growing neuroimaging databases can demonstrate systematic <u>associations</u> between specific conscious contents and the activation of specific cortical areas by using meta-analytic <u>reverse inference</u> (Poldrack, 2006; Yarkoni et al., 2010; Moran and Zaki, 2013;Poldrack and Yarkoni, 2016). Third, <u>multivariate decoding techniques</u> can compare the <u>predictive value</u> of various NCC candidates for specific conscious percepts (Haynes, 2009; Sandberg et al., 2014).

### III.1. Forward inference: Distilling the true NCC

As was argued above, the cleanest way to identify the <u>full NCC</u> is to use within-state, no-task paradigms (Fig. 3A-D), which avoid confounds due to behavioral state changes and dissociate consciousness from behavioral responsiveness and task performance. Within-state studies contrasting dreaming versus non-dreaming during NREM sleep and REM sleep, have pointed to a ‘posterior hot zone’ in parieto-occipital areas, possibly extending to mid-cingulate regions, as a reliable NCC (Siclari et al., 2017). Within-state contrasts applied to brain damaged patients, comparing VS to MCS, also reveal most consistent differences within the posterior cortex (Vanhaudenhuyse et al., 2010; Wu et al., 2015).

**Fig. 3.**
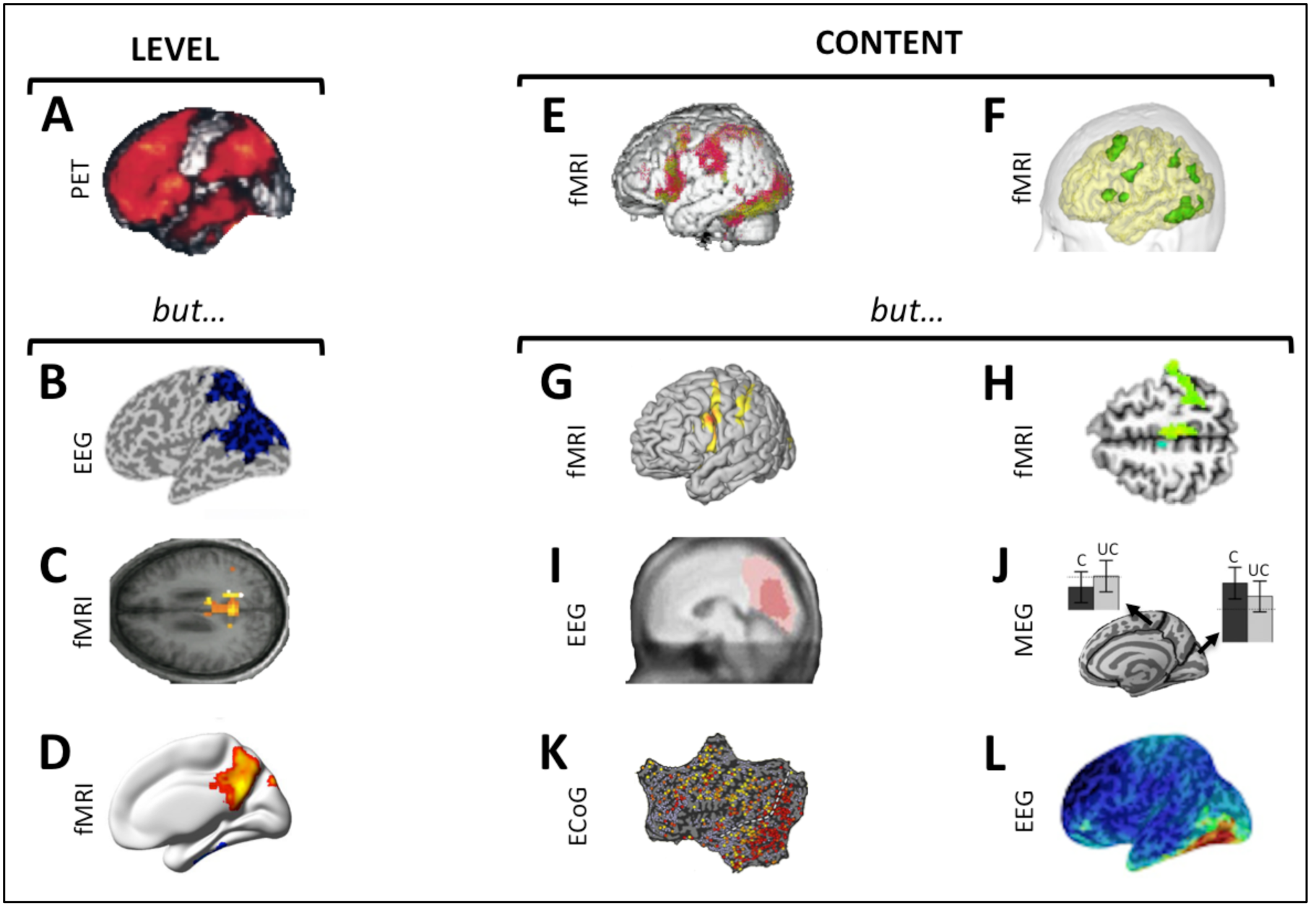
Neuroimaging: forward inference. *Level of consciousness.* **(A)** Between-state paradigm contrasting brain activity during NREM sleep and wakefulness (Kajimura et al., 1999) shows a relative deactivation of fronto-parietal cortices. **(B)** When subjects are awoken from NREM sleep and asked if they experienced anything prior to being awakened, EEG data during dream experiences shows reduced low-frequency activity (1-10 Hz) compared to dreamless sleep in a posterior parieto-occiptal hot zone (Siclari et al., 2014). **(C-D)** Comparing patients in a minimum conscious state (MCS) to patients in a vegetative state (VS) reveals differences in connectivity restricted to posterior cortex (C: (Vanhaudenhuyse et al., 2010); D: (Wu et al., 2015)). *Content of consciousness.* **(E-F)** Tasks involving reporting seen versus unseen stimuli highlight differences in fronto-parietal cortices (E: binocular rivalry (Lumer et al., 1998); F: visual word masking tasks (Dehaene et al., 2001)). (G) When conscious visual perception is dissociated from behavior (*i.e.* button press), only differences in activity in occipital and parietal regions remain (Frässle et al., 2014). **(H)** Conscious perception of weak somatosensory stimuli correlate with cortical changes in BOLD signal restricted to contralateral rolandic and parietal areas (Meador et al., 2017); **(I-J)** An early ‘visual awareness negativity’ around 200 msec in posterior temporal and occipital areas is found in two masking paradigms (I: (Koivisto and Revonsuo, 2010); J: (Andersen et al., 2016); C: conscious stimulus; UC: unconscious stimulus). **(K)** Visual one-back paradigm in patients implanted with subdural electrode arrays. The visual cortex (right of the dashed white line) responds rapidly to the seen stimulus (red) while frontal regions are modulated by the task (yellow) (Noy et al., 2015). **(L)** A within-state no-task experiment (Fig. 1D) contrasting EEG activity during dreams with and without faces, highlighted the fusiform gyrus as content-specific NCC (Siclari et al., 2014).

With respect to the <u>content of consciousness</u> (Fig. 3E-L), the use of stimuli that are task irrelevant makes it possible to dissociate between the true NCC and various cognitive functions involved in behavioral demands (Aru et al., 2012; de Graaf et al., 2012). During both inattentional blindness (Pitts et al., 2012) and backward masking (Pitts et al., 2014) paradigms, the NCC for task-irrelevant percepts are located in posterior cortex, while a difference in frontal activity (P3 potential) is only present if stimuli are task-relevant. ‘No-report’ paradigms have also pointed to posterior regions as the NCC, while frontal cortex activation is correlated with reporting (Tsuchiya et al., 2015; Koch et al., 2016b). Similar dissociations have been identified by orthogonally manipulating consciousness versus selective attention (Koch and Tsuchiya, 2012), working memory (King et al., 2016) or expectation (Melloni et al., 2011). During REM sleep, a ‘no-task’ state, specific dream contents such as faces, movement, speech or spatial aspects can be predicted from posterior, but not anterior cortex (Siclari et al., 2017). The same approach has highlighted a potential contribution of mid-cingulate cortex to conscious thought, whether during waking, NREM and REM sleep (Perogamvros et al., 2017; Siclari et al., 2017). Finally, when meditation practitioners become immersed in a state of vivid imagery, activity in their frontal lobe decreases (Lou and colleagues 1999).

### III.2. Reverse inference: content-specific NCC versus content-unspecific cognitive processes

To systematically assess the lawfulness of NCC candidates, meta-analyses should pool evidence across a large number of experiments. In recent years, several open-access frameworks have been developed to pool data from thousands of neuroimaging studies (Eickhoff et al., 2011; Poldrack and Yarkoni, 2016). For example, the Neurosynth framework (http://www.neurosynth.org) (Yarkoni et al., 2011) combines an automated tool to extract activation coordinates with a taxonomy of cognitive processes that includes a few entries for consciousness. Such tools should be used cautiously, because their constituent neural data consist of 3D coordinates of reported peaks of activity, rather than unthresholded statistical maps (but see Neurovalt, http://www.neurovault.org. (Gorgolewski et al., 2016)) and they rely on functional labels assigned by investigators. Nonetheless, these meta-analytic tools can already be useful to illustrate reverse inference and to generate hypotheses.

When using Neurosynth in a traditional meta-analysis approach - computing the probability that different brain regions are active when the study topics include consciousness – activation maps display some parts of the frontal cortex (see Fig. 4A). However, when using Neurosynth for reverse inference – computing the probability that consciousness is mentioned in the study given the activation of different brain regions - frontal cortex activation disappears (Fig. 4B). By contrast, in agreement with lesion and stimulation studies, reverse inference locates the best predictor of face percepts in the right FFA (Fig.4C-D). Content-specific results for visual words or motion, speech sounds, or touch perception are likewise regionalized to occipital, temporal and parietal cortices. In all these cases, reverse inference does not highlight frontal areas as predictive for specific conscious contents.

**Fig. 4.**
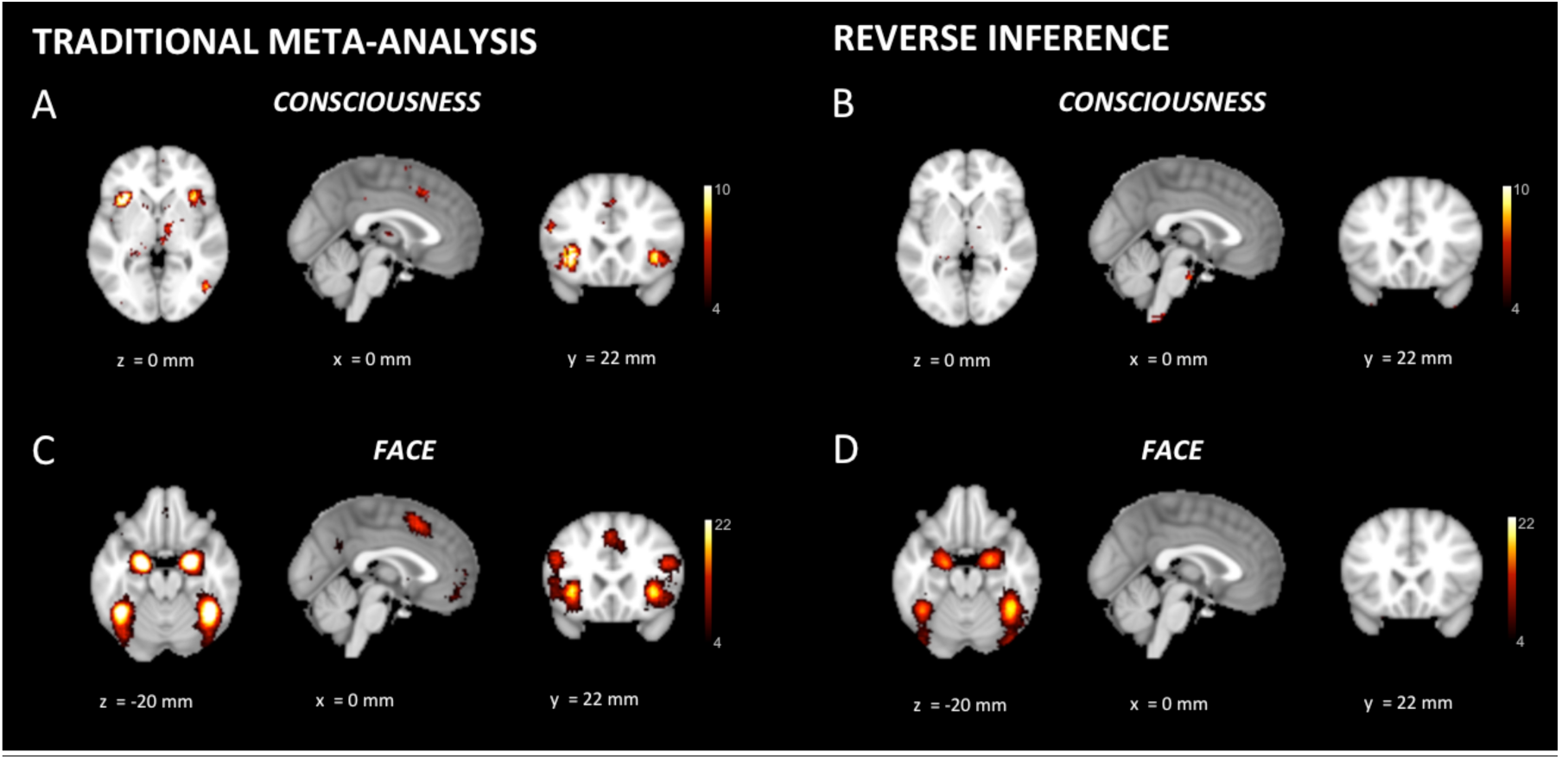
Neuroimaging: reverse inference. **(A)** When using Neurosynth for a traditional ‘forward’ meta-analysis - computing the probability that different brain regions are active when the topics of a study include consciousness - parts of frontal cortex show up. **(B)** When using Neurosynth for reverse inference - computing the probability that consciousness is included within the topics of a study, given the activation of different parts of the brain - frontal cortex disappears. The key term ‘conscious’ was used on the Neurosynth website to extract both ‘forward’ metaanalysis and reverse inference analysis steps in A-B. **(C)** The same frontal areas that identified in a ‘forward’ meta-analysis for consciousness also appear activated in a ‘forward’ meta-analysis for faces. **(D)** In contrast, reverse inference for faces no longer identifies frontal cortex activity, but rather locates the activation predicting highest probability for face percepts in the right FFA. The key term ‘faces’ was used on the Neurosynth website to extract both ‘forward’ meta-analysis and reverse inference analysis steps in C-D. x,y,z values represent MNI coordinates, and a color scale is used for Z values.

Neurosynth also permits the assessment of the functional specificity of brain areas at user-specified coordinates. For example, activation in the FFA (with coordinate selected from the traditional ‘forward’ meta-analysis, [41 −49 −20]) is consistently predictive for faces (p = 0.88), temporo-occipital cortex for visual words ([−46 −54 - 12], p = 0.86) or visual motion ([46 −68 0], p =0.9), superior temporal cortex for speech sounds ([−58 −10 0], p =0.84), and postcentral cortex for touch ([−54 −22 20], p = 0.88). In contrast, the statistical maximum of frontal cortex activation obtained from the ‘traditional meta-analysis on consciousness ([−47 6 28]) is found to be most predictive for the terms ‘phonological’ and ‘task’ (p = 0.76 and p = 0.63, respectively).

### III.3. Prospective predictive approaches: decoding consciousness in individual trials/subjects

Ideally, decoding approaches would identify the true NCC as neural activity patterns most predictive for the presence of a given conscious content (Sandberg et al., 2014). Unlike classical statistical analysis, decoding approaches also assess reproducibility as the percentage of accurate classification among single trials (Haynes, 2009). With respect to the <u>full NCC,</u> the best predictors for differentiating MCS from VS using positron emission tomography (PET) or fMRI were located in parietal, temporal and occipital cortices ((Demertzi et al., 2015; Stender et al., 2016); Fig. 5, Upper Panel). An online prospective approach based on EEG markers of arousal in posterior cortex was able to predict consciousness (dreaming) versus unconsciousness during NREM sleep with 85% accuracy (Siclari et al., 2017). *Post-hoc* analysis also located the areas most predictive for dreaming consciousness within temporo-parieto-occipital cortices (Siclari et al., 2017). Anesthesia studies have shown that frontal activity is a poor predictor of consciousness (Avidan et al., 2011), but there are so far no data for posterior cortex.

**Fig. 5.**
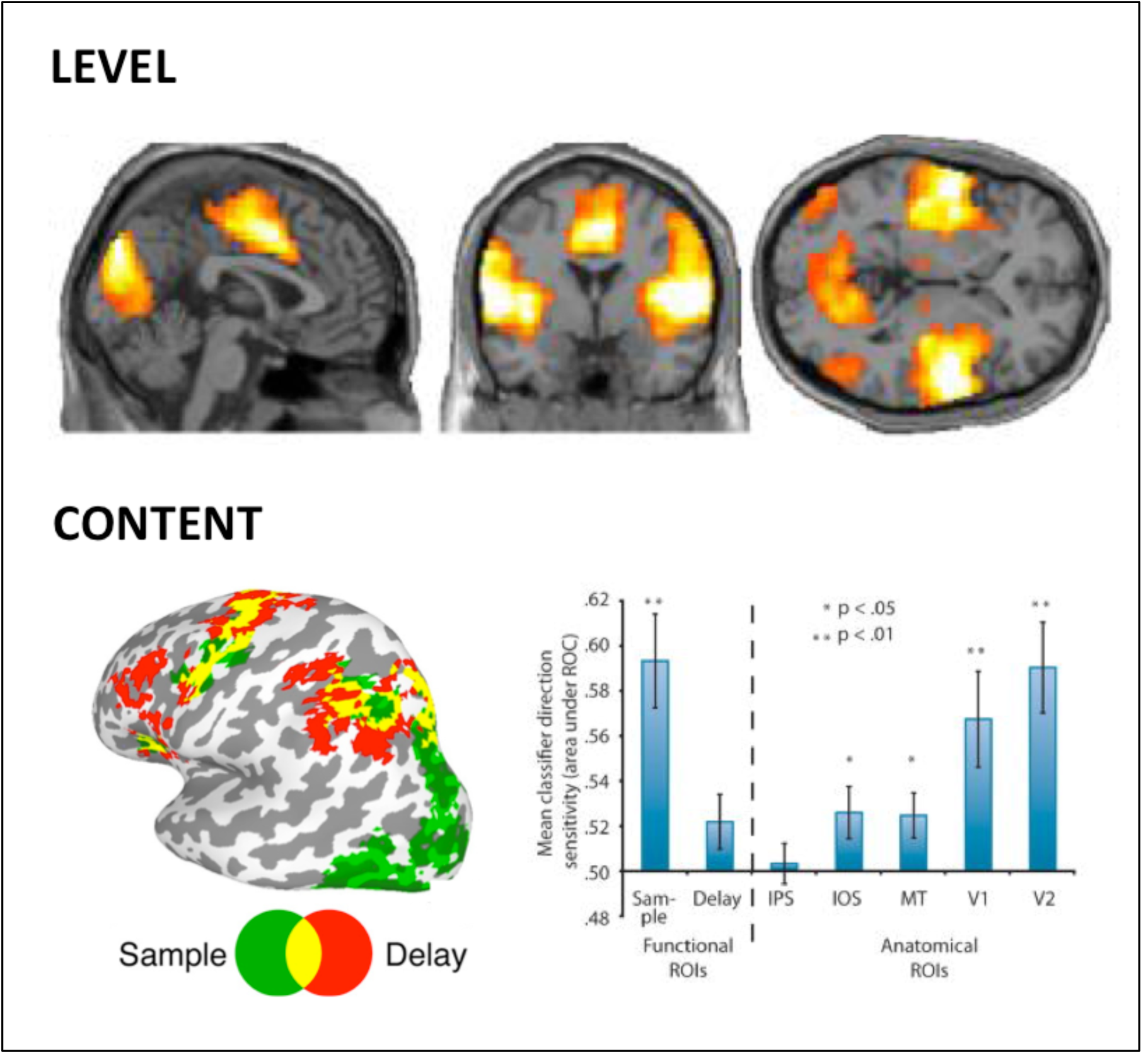
Neuroimaging: predictive approaches. Upper panel: machine learning approaches applied to functional MRI resting state show that temporo-parieto-occipital connectivity best differentiates patients in MCS versus VS (Demertzi et al., 2015). Lower panel: the contents of a working memory task can best be decoded from the back of the brain (green ROI), but not from the front of the brain (red ROI) (Emrich et al., 2013). Right side panel shows classification accuracy compared to chance to identify conscious contents from both ROIs and from different anatomical locations across the visual hierarchy. IPS: intra-parietal sulcus; IOS: intra-occipital sulcus; MT: area MT; V1: primary visual cortex; V2: secondary visual cortex; ROI: region of interest.

As for <u>content-specific NCC,</u> numerous studies in both awake and dreaming subjects could decode the occurrence of specific experiential contents from the activity of specific regions of posterior cortex (Nishimoto et al., 2011; Horikawa et al., 2013; Siclari et al., 2017). Working memory contents can also be decoded more reliably from the back than from the front of the cortex ((Emrich et al., 2013); Fig. 5, Lower Panel). Finally, multivariate patterns predictive of differences in subjective experiences both within-(Kriegeskorte, 2011) and between subjects (Charest et al., 2014) are most consistently found in posterior cortex.

## IV. Conclusion and future directions

In this Perspective, we reviewed evidence across lesion, stimulation and recording studies that consistently point to the ‘back’ of the cortex as a ‘posterior hot zone’ playing a direct role in specifying the contents of consciousness. By contrast, evidence for a direct involvement of the ‘front’ of the cortex is missing or unclear. At a minimum, reports of conscious patients after bilateral frontal lobectomy demonstrate that the frontal cortex is not necessary for consciousness. While most prefrontal regions may be “mute” as regards to consciousness, not unlike basal ganglia and cerebellum, it remains possible that some prefrontal regions – such as ventromedial areas (Koenigs et al., 2007) - contribute specific conscious contents such as feelings of reflection, valuation, and affect (Koch et al., 2016a). Below we discuss promising future research directions.

***Lesion studies*** would benefit from a systematic assessment of loss of specific conscious contents after frontal cortex damage, sampling both task-related experiences as well as dream contents (as in (Solms, 2014)). Future experiments should also investigate possible dissociations between consciousness and cognitive functions after frontal damage, detail the precise 3D location (as in (Mah et al., 2014)) and laminar profile (Koch et al., 2016b) of the lesions, and control for network effects (Fischer et al., 2016).

***Stimulation studies*** should explore the effects of local perturbations on both task performance (as in (Winawer and Parvizi, 2016)) and subjective experience (using structured questionnaires). Direct electrical stimulation combined with intracranial recordings at the stimulation site and at distant sites (as in (Keller et al., 2014; Pigorini et al., 2015)) should help to identify specific patterns of functional connectivity involved in consciousness.

***Neuroimaging studies*** should further exploit within-state, no-task paradigms to differentiate between the full NCC and neural correlates of responsiveness (Koch et al., 2016b). With respect to conscious content, pooling across an exhaustive set of different experiments (as in (Axelrod et al., 2015)) - including a combination of report and no-report paradigms (Tsuchiya et al., 2016) - may help to identify content-specific NCC as the brain regions most consistently activated in the presence of specific conscious percepts. Once several content-specific NCC are identified, it will become meaningful to carry out a formal comparison between them (Rutiku et al., 2016). Systematic meta-analyses using reverse inference will be useful to assess the lawfulness of NCC candidates while avoiding cherry-picking (Moran and Zaki, 2013). Meta-analytic frameworks such as Neurosynth should be modified by explicitly incorporating the dissociation between consciousness and cognitive functions such as attention and task execution. Prospective studies should confirm that the candidate NCC identified through forward and reverse inferences remain the best predictors for the presence of consciousness across different physiological or pathological states, at the level of single trials, or even online (as in (Siclari et al., 2017)). Decoding studies should also explicitly compare the predictive value of different neural activity patterns for specific conscious contents (as in (Emrich et al., 2013)). Finally, prospective studies should be used to assess the clinical utility of different NCC candidates for detecting consciousness in brain damaged patients (as in (Demertzi et al., 2015; Stender et al., 2016)).

## Acknowledgments

This work was supported by NIH/NINDS 1R03NS096379 (to M.B.), by the Tiny Blue Dot Foundation and the Distinguished Chair in Consciousness Science (University of Wisconsin) (to G.T.), and by the James S. McDonnell Scholar Award 2013 (to M.M.).

## Notes

**Conflicts of Interest**: The authors declare no conflicts of interest.

